# Global biogeography of mangrove sediment microbiomes is structured by a conserved core and environmental selection

**DOI:** 10.64898/2026.01.07.698013

**Authors:** Francisco J. Balvino-Olvera, Mariana Pamela Mota-Montes de Oca, Daniel Bustos-Díaz, Nelly Sélem-Mojica, DJ Jimenez, Irma González-González, Pindaro Diaz, Mirna Vázquez-Rosas-Landa

**Affiliations:** Unidad Académica de Ecología y Biodiversidad Acuática, Instituto de Ciencias del Mar y Limnología, Universidad Nacional Autónoma de México, Ciudad de México, México; Centro de Ciencias de Matemáticas, Antigua Carretera a Pátzcuaro 8701 Col. Ex Hacienda de San José de la Huerta, C.P. 58089 Morelia, Michoacán, México; Biological and Environmental Sciences and Engineering Division (BESE), King Abdullah University of Science and Technology (KAUST), Thuwal, Kingdom of Saudi Arabia

## Abstract

Mangrove sediments are globally important biogeochemical hotspots, yet the large-scale organization and assembly of their microbiomes remain poorly resolved. Here, we compiled and analysed 390 shotgun metagenomes from mangrove sediments across 43 sites worldwide to quantify how diversity, spatial turnover, and ecological processes structure microbial communities across environmental and geographic gradients. We identify a striking dual architecture in mangrove sediment microbiomes. A remarkably small and ubiquitous taxonomic core, representing a minor fraction of total richness but dominating community abundance, persists across continents, climatic regimes, and marine realms. This core is composed primarily of anaerobic microbial lineages associated with carbon, nitrogen, and sulfur cycling and shows limited spatial and environmental turnover. In contrast, the non-core fraction is highly diverse, responds strongly to climatic and edaphic gradients, and drives most biogeographic differentiation. Assembly-process modelling reveals that deterministic selection dominates global taxonomic turnover, with homogeneous selection stabilizing shared community components and heterogeneous selection promoting regional differentiation, while dispersal limitation plays a secondary role. Network analyses further show that core taxa occupy structurally central positions in microbial co-occurrence networks, supporting overall connectivity despite pronounced regional variation in peripheral community composition. Together, these results demonstrate that mangrove sediment microbiomes combine a conserved functional backbone with an environmentally responsive periphery. This organization reconciles global functional continuity with strong regional differentiation and provides a basis for anticipating microbiome responses to environmental change in mangrove ecosystems.

## Introduction

Mangrove ecosystems are among the most productive coastal environments on Earth and play a major role in global carbon cycling. Their sediments accumulate large amounts of organic matter, sustain intense anaerobic metabolism, and act as long-term carbon sinks that mitigate atmospheric greenhouse gas emissions ^1,2^. These processes are largely mediated by complex microbial communities that drive carbon, nitrogen, and sulfur cycling under steep redox and chemical gradients^3–6^. Understanding how mangrove sediment microbiomes are structured and maintained is therefore central to predicting the stability and resilience of these ecosystems under accelerating environmental change ^2^.

Over the past decade, numerous studies have characterized mangrove-associated microbial communities across local and regional scales, revealing high taxonomic diversity and strong sensitivity to environmental gradients such as salinity, nutrient availability, and redox conditions ^3–6,7^. Most of this work has relied on amplicon-based surveys, which have been instrumental in identifying dominant microbial groups and documenting patterns of spatial turnover^8–10^. However, these approaches provide limited insight into the functional potential of communities and into the ecological processes that govern their assembly. As a result, despite growing datasets, our understanding of how mangrove sediment microbiomes are organized at the global scale remains fragmentary and largely taxonomic^11–13^.

In particular, three fundamental questions remain unresolved. First, it is unclear whether mangrove sediment microbiomes exhibit consistent large-scale biogeographic structure or are primarily shaped by local environmental conditions ^7,8,14^. Second, the relative roles of environmental selection, dispersal limitation, and stochastic processes in assembling these communities across continents have not been quantified explicitly^15,16^. In microbial community ecology, deterministic selection refers to reproducible, environment-driven differences in community composition, whereas stochastic processes reflect random demographic variation and dispersal. Third, it remains unknown whether a globally conserved microbial core underpins mangrove sediment microbiomes, and if so, how such a core coexists with strong regional differentiation^17,18^.

Here, we address these questions by compiling and analysing 390 shotgun metagenomes from mangrove sediments spanning 43 sites worldwide. By integrating taxonomic, environmental, spatial, and network-based analyses, we quantify global patterns of diversity and turnover, identify the ecological processes driving community assembly, and assess the structure and functional significance of core and non-core community fractions. Our approach builds on quantitative frameworks developed to disentangle microbial assembly processes across ecosystems^19^. It extends previous amplicon-based biogeographic syntheses^8,10^ by explicitly incorporating functional potential, spatial structure, and network topology at the global scale.

Our results reveal that mangrove sediment microbiomes are organized around a dual architecture, in which a small, globally conserved core provides functional continuity, while a large and environmentally responsive periphery drives biogeographic differentiation. This framework reconciles global stability with regional variability and offers a basis for predicting microbiome responses to environmental change and for incorporating microbial processes into mangrove conservation and restoration strategies ^1,2,20^.

## RESULTS AND DISCUSSION

### Global patterns of alpha diversity in mangrove sediments

Mangrove-sediment microbiomes exhibited high alpha diversity across global scales, with marked geographic structure. After rarefaction, Hill diversity (q = 1) varied substantially among marine biogeographic realms, with the highest values observed in Central and Western Indo-Pacific mangroves and consistently lower diversity in the Tropical Atlantic. Spatial modelling using generalized additive models (GAMs) revealed a pronounced east–west gradient, identifying the Indo-Pacific as a global hotspot of microbial diversity in mangrove sediments (Fig. 1A).

**Figure 1.**
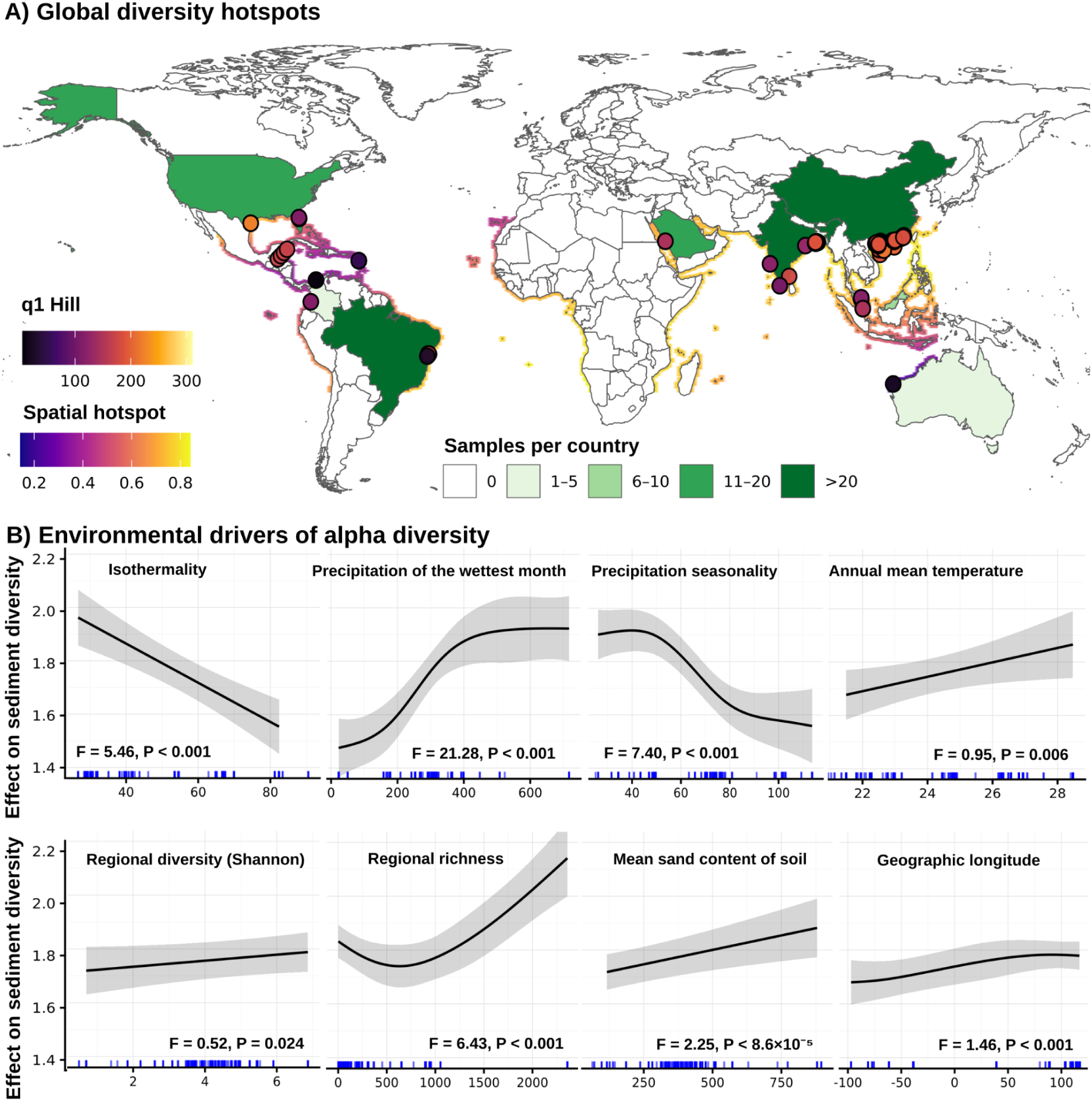
Global patterns and environmental drivers of alpha diversity in mangrove-sediment microbiomes. **(A)** Spatial hotspots of microbial alpha diversity (Hill q = 1) across tropical and subtropical mangroves. Colours show the predicted diversity surface from a generalized additive model (GAM) fitted to log-transformed Hill q = 1 with a two-dimensional spatial smooth (latitude-longitude); warmer colours indicate higher expected diversity. Circles mark sampling sites, and outline colours indicate the number of samples per country. **(B)** Partial effects of key environmental predictors on mangrove-sediment diversity from the same GAM. Curves show the smooth effect of each predictor on log-transformed Hill q = 1 (centred on the mean effect), with shaded bands representing 95% confidence (± 2 se). Predictors are: isothermality (BIO3), precipitation of the wettest month (BIO13), precipitation seasonality (BIO15), annual mean temperature (BIO1), regional Shannon diversity (GBIF Shannon), regional species richness (GBIF richness), mean sand content of surface soil (sand_mean), and geographic longitude (lon). For each smooth, the corresponding F statistic and P value from the GAM are shown inside the panel.

Climatic variables emerged as the dominant drivers of alpha diversity. Precipitation during the wettest month exerted the strongest effect, with diversity increasing sharply from dry conditions and stabilizing at high rainfall levels. This response was modulated by climatic variability: the most diverse communities occurred under low precipitation seasonality and low isothermality, indicating that sustained water availability combined with seasonal thermal variation promotes microbial niche diversification (Fig. 1B). Mean annual temperature showed a weaker but positive effect, suggesting that warmer mangroves tend to host slightly more diverse microbiomes when moisture conditions are favourable.

Beyond climate, sediment texture, and regional biodiversity context further shaped diversity patterns. Alpha diversity increased with mean sand content, consistent with coarser sediments generating steeper redox and chemical microgradients that can be exploited by distinct microbial guilds ^21^. In addition, microbial diversity was positively associated with regional macro-organism richness and Shannon diversity derived from GBIF, indicating that above and below-ground biodiversity are coupled across biogeographic gradients ^22^.

Together, these results show that global patterns of mangrove-sediment microbial diversity are primarily structured by climatic water availability and variability, with sediment properties and regional biodiversity modulating local richness. The Indo-Pacific diversity hotspot thus reflects not only historical biogeography, but also a convergence of hydrological, climatic, and sedimentary conditions that maximize microbial niche availability at the global scale, consistent with recent mangrove sediment syntheses ^8,10^.

### Distance–decay and regional structuring of mangrove-sediment microbiomes

To characterize large-scale spatial patterns in mangrove-sediment microbiomes, we quantified distance–decay relationships across global spatial scales. Community similarity declined steadily with increasing geographic distance, from co-located sites to pairs separated by nearly 20,000 km, indicating that geographically proximate mangroves host more similar microbial assemblages than distant ones (Fig. 2A). This pattern was evident for both abundance-based (Bray–Curtis) and presence-absence (Sørensen) metrics.

**Figure 2.**
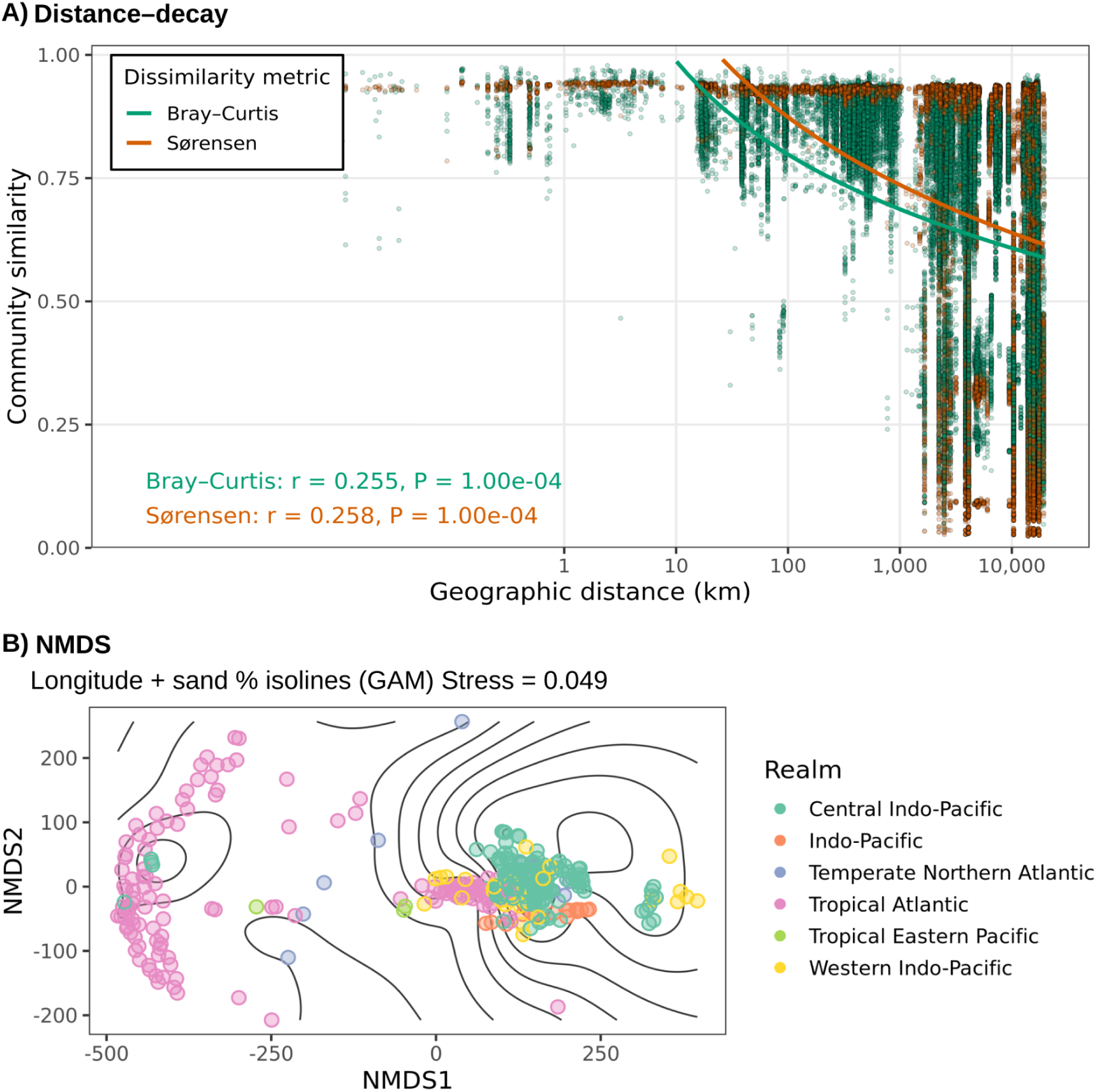
Spatial structuring of mangrove-sediment beta diversity. **(A)** Distance-decay of community similarity across global mangrove sediments. Points show pairwise community similarity (1 − dissimilarity) as a function of geographic distance between sites (log₁₀ km) for abundance-based (Bray-Curtis, green) and presence-absence (Sørensen, orange) metrics. Solid lines are linear fits for each metric. Both dissimilarities increased significantly with distance (Mantel tests, Bray-Curtis: r = 0.255, P = 1.0×10⁻⁴; Sørensen: r = 0.258, P = 1.0×10⁻⁴; 9,999 permutations), and the slopes differed between metrics (permutational ANCOVA interaction: F = 233.39, P = 0.001), indicating stronger spatial turnover in relative abundances than in mere species presence-absence. **(B)** Non-metric multidimensional scaling (NMDS) of community composition based on Aitchison distances, with generalized additive model (GAM) isolines for geographic longitude. Points represent sites and are coloured by marine realm; isolines depict predicted longitude values in NMDS space. The ordination (stress = 0.049) reveals a clear compositional gradient linked to both longitude and biogeographic realm, with Indo-Pacific realms clustering together and Atlantic mangroves occupying a distinct portion of ordination space.

However, the rate of decay differed between metrics. Abundance-weighted dissimilarity increased with distance more rapidly than presence-absence dissimilarity, indicating that many taxa are shared across distant mangroves but undergo substantial reshuffling in relative dominance along spatial and environmental gradients. This contrast suggests that widespread taxa persist globally, while changes in abundance structure contribute disproportionately to spatial differentiation, consistent with the coexistence of cosmopolitan and regionally structured community fractions described in other microbial systems ^23^.

Ordination analyses further revealed a strong regional organization of community composition. Non-metric multidimensional scaling (NMDS) based on Aitchison distances showed a pronounced east-west gradient, with Indo-Pacific and Atlantic mangroves occupying largely distinct regions of ordination space (Fig. 2B). Longitude closely tracked this compositional gradient, indicating that broad-scale geographic structure aligns with underlying environmental gradients distributed across ocean basins, consistent with previous mangrove sediment biogeographic syntheses ^6,8,10^.

Permutation-based analyses confirmed that compositional differences among mangrove-sediment microbiomes are structured across multiple spatial scales. Countries and ecoregions explained a substantial fraction of community variation, while marine realms accounted for a smaller but still significant component. This pattern indicates that although major ocean basins impose a biogeographic scaffold, much of the differentiation in mangrove-sediment microbiomes emerges at regional to subregional scales, where climate, sediment properties, vegetation, and local disturbance regimes vary most strongly, as reported in regional studies of mangrove sediments and soils ^7,24^.

Together, these results demonstrate that mangrove-sediment microbiomes are strongly spatially structured across global scales, with pronounced turnover in community composition that cannot be explained by geographic distance alone. ^15,16^.

While distance–decay and ordination analyses reveal strong spatial structuring of mangrove-sediment microbiomes, they do not identify the environmental and spatial factors underlying this structure. We therefore used multivariate and variance-partitioning approaches to disentangle the relative contributions of environmental gradients and spatial structure to global beta diversity.

### Environmental and spatial drivers of beta diversity

To identify the environmental and spatial factors underlying global patterns of beta diversity, we analysed mangrove-sediment microbiomes using distance-based redundancy analysis (db-RDA) and variance partitioning. Distance-based redundancy analysis (db-RDA) constrained by climatic, edaphic, macroecological, and anthropogenic variables revealed a strong compositional gradient aligned primarily with precipitation, temperature, regional biodiversity context, and sediment properties (Fig. 3A). Together, these variables captured the main axes of taxonomic turnover across geographically distant mangroves.

**Figure 3.**
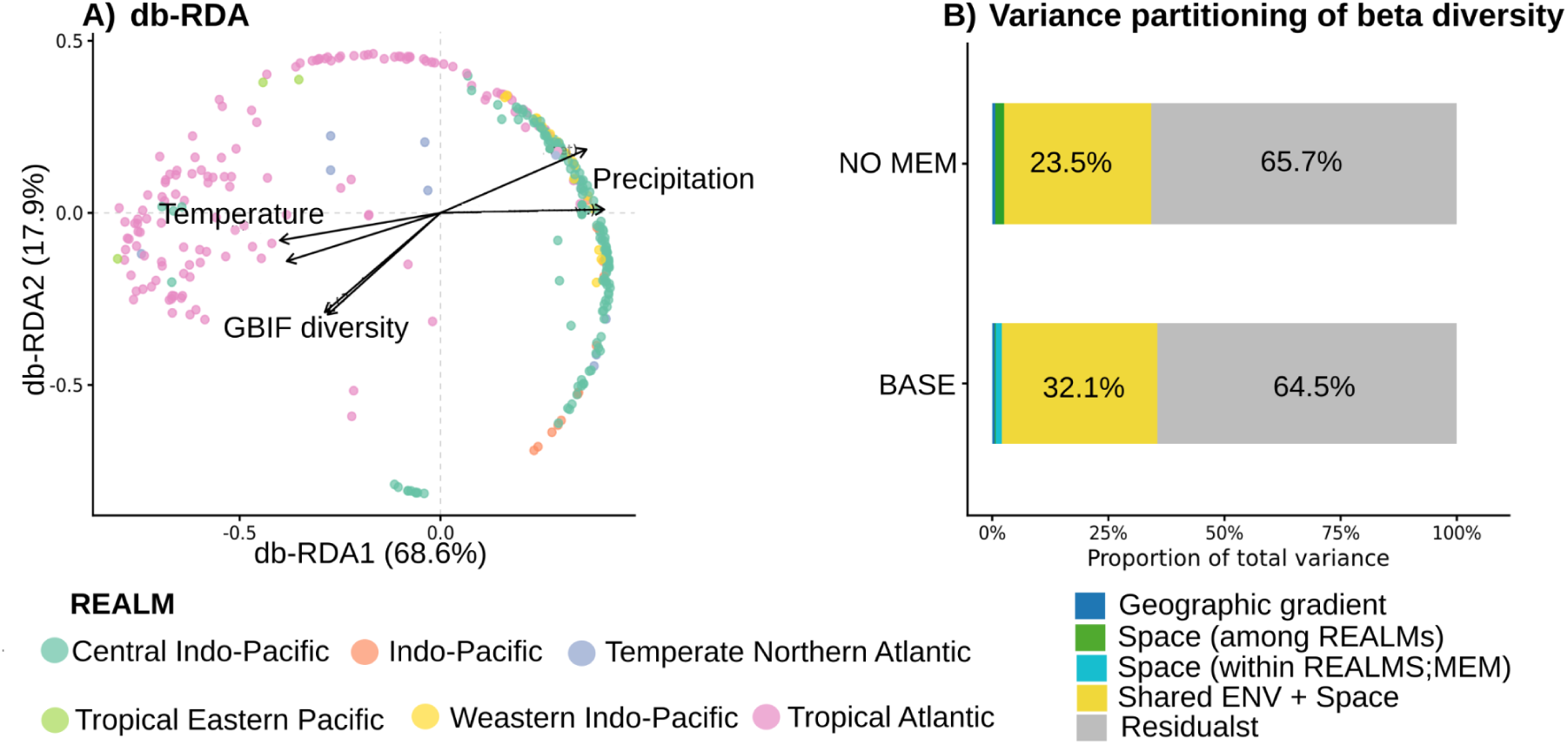
Environmental and spatial controls on mangrove-sediment beta diversity. **(A)** Environment-constrained db-RDA of Bray-Curtis community dissimilarities (n = 329). Points are sites coloured by marine realm; arrows are the most significant environmental vectors from envfit (999 permutations), including precipitation (BIO13, BIO15), temperature (BIO1-BIO3), regional GBIF richness/Shannon, and soil properties. The first two constrained axes explain 68.6% and 17.9% of the constrained variance, respectively. **(B)** Variance partitioning from db-RDA comparing two specifications: BASE (ENV + lat/lon + realm + within-realm spatial eigenfunctions, MEM) and NO MEM (ENV + lat/lon + realm). Stacked bars show the fraction of total variance attributed to the geographic gradient (lat/lon), space among realms, space within realms (MEM; BASE only), the shared ENV + space component, and residuals. Global adjusted R²: 0.356 (BASE) and 0.343 (NO MEM). In BASE, most explained variance is shared ENV + space (32.1%), with small pure fractions for environment (1.4%), geographic gradient (0.5%), and MEM (1.3%), and a negligible, non-significant pure realm effect (0.2%). Omitting MEM reallocates variance from the shared component to pure environment (8.2%) and pure realm (1.8%), while the geographic gradient remains minor (0.7%).

Variance partitioning clarified how environmental and spatial factors interact to structure this turnover. In the full model, most of the explained variation was attributed to the shared fraction between environment and space, whereas the pure effects of environment, large-scale geographic gradients, and marine realms were individually small (Fig. 3B). This pattern indicates that microbial communities primarily track environmental gradients that are themselves spatially autocorrelated, rather than responding independently to geography or biogeographic region, consistent with expectations when environmental selection acts along spatially structured gradients ^15,25^.

When fine-scale spatial structure within realms was not explicitly modelled, the apparent contribution of pure environmental effects increased, while the influence of realms also appeared stronger. This shift demonstrates that analyses that ignore spatial autocorrelation can overestimate the independence of environmental and regional effects. By accounting for spatial structure at multiple scales, our results show that whether a mangrove is located in the Indo-Pacific or Atlantic explains little additional variation once climate, sediment properties, and regional biodiversity are considered, consistent with regional work showing strong environmental control of mangrove sediment community composition ^6–8,24^.

Overall, these analyses indicate that environmental selection acting along spatially structured gradients dominates global patterns of mangrove-sediment beta diversity. Geographic distance and regional history impose a secondary scaffold, but do not override the influence of climate, sediment chemistry, and ecosystem context ^15,16^.

### Community assembly processes reveal dominance of selection

To move from patterns of beta diversity to the mechanisms underlying community turnover, we quantified the relative contributions of selection, dispersal, and stochasticity using phylogenetically informed null models^19^. Across global scales, deterministic selection overwhelmingly dominated taxonomic turnover, accounting for most pairwise community differences. Both homogeneous and heterogeneous selection contributed substantially, indicating that strong environmental filtering promotes convergence under similar conditions while enabling divergence across contrasting local environments.

Homogeneous selection was most prevalent among communities occupying similar climatic and sedimentary environments, consistent with the persistence of shared taxa across distant regions. In contrast, heterogeneous selection increased sharply across environmental gradients, particularly those associated with precipitation, temperature variability, and sediment properties. This balance between stabilizing and diversifying selection explains how mangrove-sediment microbiomes maintain continuity in core functions while generating pronounced regional differentiation, consistent with metacommunity and niche-stochasticity frameworks ^16,26^.

Dispersal limitation contributed a smaller but consistent fraction of community turnover, particularly at intermediate to large spatial scales. This effect indicates that although mangrove sediments are not freely mixing microbial pools, constraints on dispersal alone cannot account for observed biogeographic patterns. Stochastic processes, including ecological drift, played a comparatively minor role across the dataset.

Together, these results demonstrate that mangrove-sediment microbiomes are assembled primarily through deterministic ecological processes acting along spatially structured environmental gradients. Selection, rather than dispersal or drift, provides the mechanistic link between global environmental heterogeneity and the spatial organization of microbial communities observed across mangrove ecosystems.

### A small global core underpins mangrove sediment microbiomes

Despite pronounced biogeographic variation in community composition, mangrove-sediment microbiomes shared a remarkably small and ubiquitous taxonomic core. This core comprised only a minor fraction of total taxonomic richness, yet accounted for the vast majority of sequencing reads across all sampled sites and marine realms (Fig. 4A). Core taxa were consistently detected in sediments spanning contrasting climatic regimes, sediment textures, and geographic regions, indicating a globally conserved microbial backbone, consistent with core-periphery structures described in other microbiomes ^17,18^.

**Figure 4.**
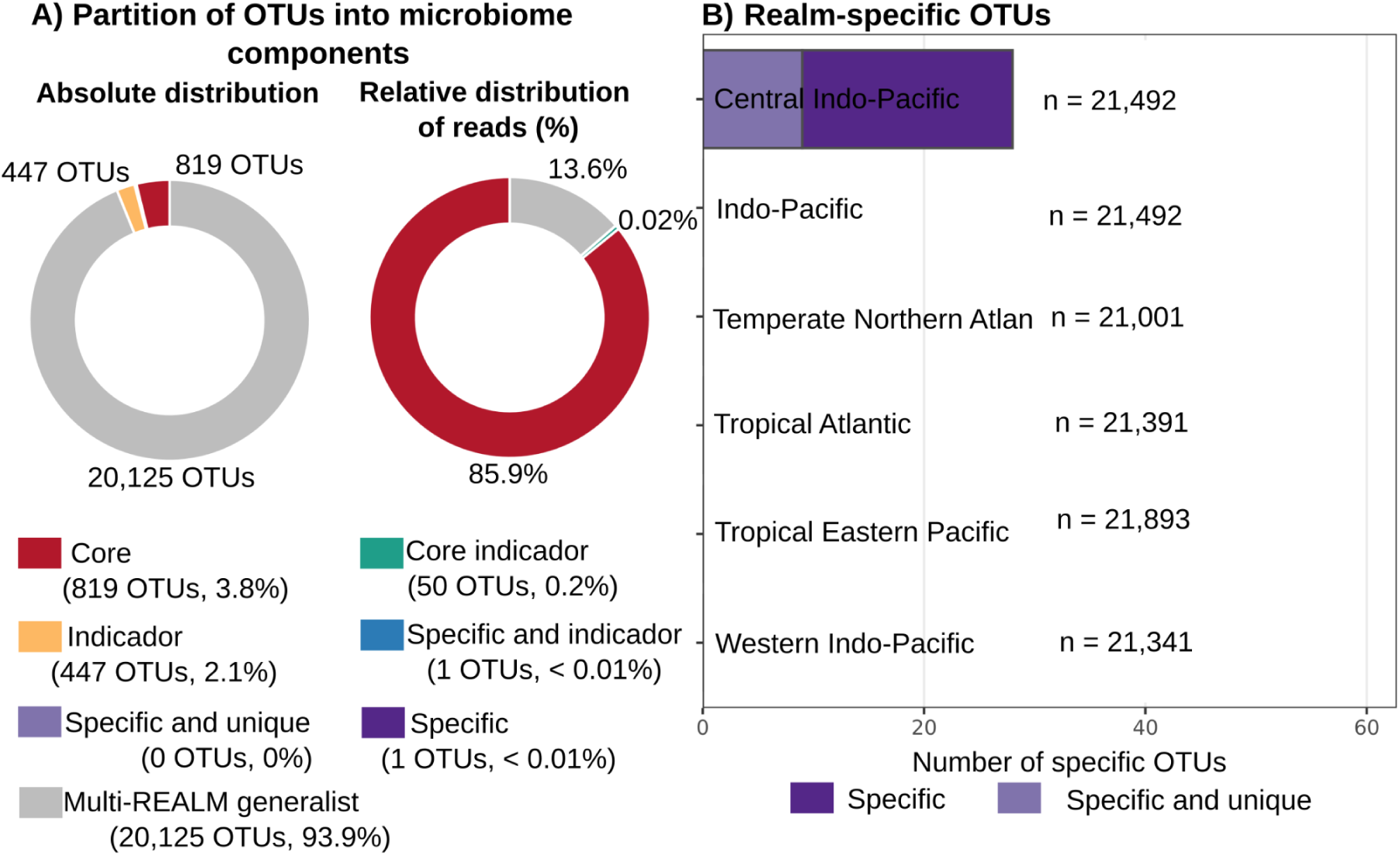
Structure and biogeographic specificity of the core mangrove-sediment microbiome. **(A)** Partition of OTUs into microbiome components. Left, absolute number of OTUs per component; right, percentage of the total read count explained by each component. Most OTUs were classified as multi-realm generalists (20,125 OTUs; 93.9% of richness), but together they accounted for only 13.6% of all reads. In contrast, the core microbiome core plus core-and-indicator OTUs (869 OTUs; 4.1% of richness) concentrated 86% of the sequencing reads. Indicator and realm-specific components were numerically rare (2.2% of OTUs in total) and contributed < 0.1% of reads. **(B)** realm-specific OTUs. Bars show, for each marine realm, the number of OTUs restricted to that realm (“specific”, dark purple) and the subset that is also confined to a single locality within that realm (“specific and unique”, light purple). Only the Central Indo-Pacific contained realm-specific OTUs (28 in total, 9 of them unique to one locality), whereas no specific OTUs were detected in the other realms despite comparable overall richness (20,800-21,400 OTUs per realm; values on the right).

Taxonomically, the core was dominated by anaerobic bacterial lineages characteristic of reduced sediment environments, including members of Desulfobacterota, Chloroflexota, Firmicutes, and related groups. These taxa encompassed functional guilds central to carbon mineralization, sulfate reduction, methanogenesis-associated processes, and nitrogen transformations, consistent with the anoxic and carbon-rich conditions of mangrove sediments and with taxa repeatedly reported as dominant in regional mangrove sediment studies^7,8,11,12,27,28^

Core taxa exhibited distinct ecological signatures compared to the remainder of the community. They showed limited spatial turnover, weak responses to climatic and edaphic gradients, and broad environmental tolerances, consistent with the prevalence of homogeneous selection identified in assembly-process analyses. In contrast, most rare and moderately abundant taxa were absent from the core and displayed strong environmental and spatial structuring.

Together, these results indicate that mangrove-sediment microbiomes are anchored by a small set of globally distributed, environmentally tolerant taxa that dominate community abundance and underpin key biogeochemical functions. This conserved core provides continuity across regions, while allowing extensive taxonomic variability in the surrounding microbial community.

### Core and non-core fractions show contrasting biogeographic turnover

Partitioning communities into core and non-core fractions revealed strikingly different biogeographic behaviours. While the full community exhibited clear distance–decay relationships, this pattern was almost entirely driven by the non-core fraction (Fig. 5). Core taxa showed weak or negligible distance–decay, maintaining similar composition and relative abundance across geographically distant mangroves. In contrast, non-core taxa displayed rapid declines in similarity with increasing geographic distance, indicating strong spatial turnover.

**Figure 5.**
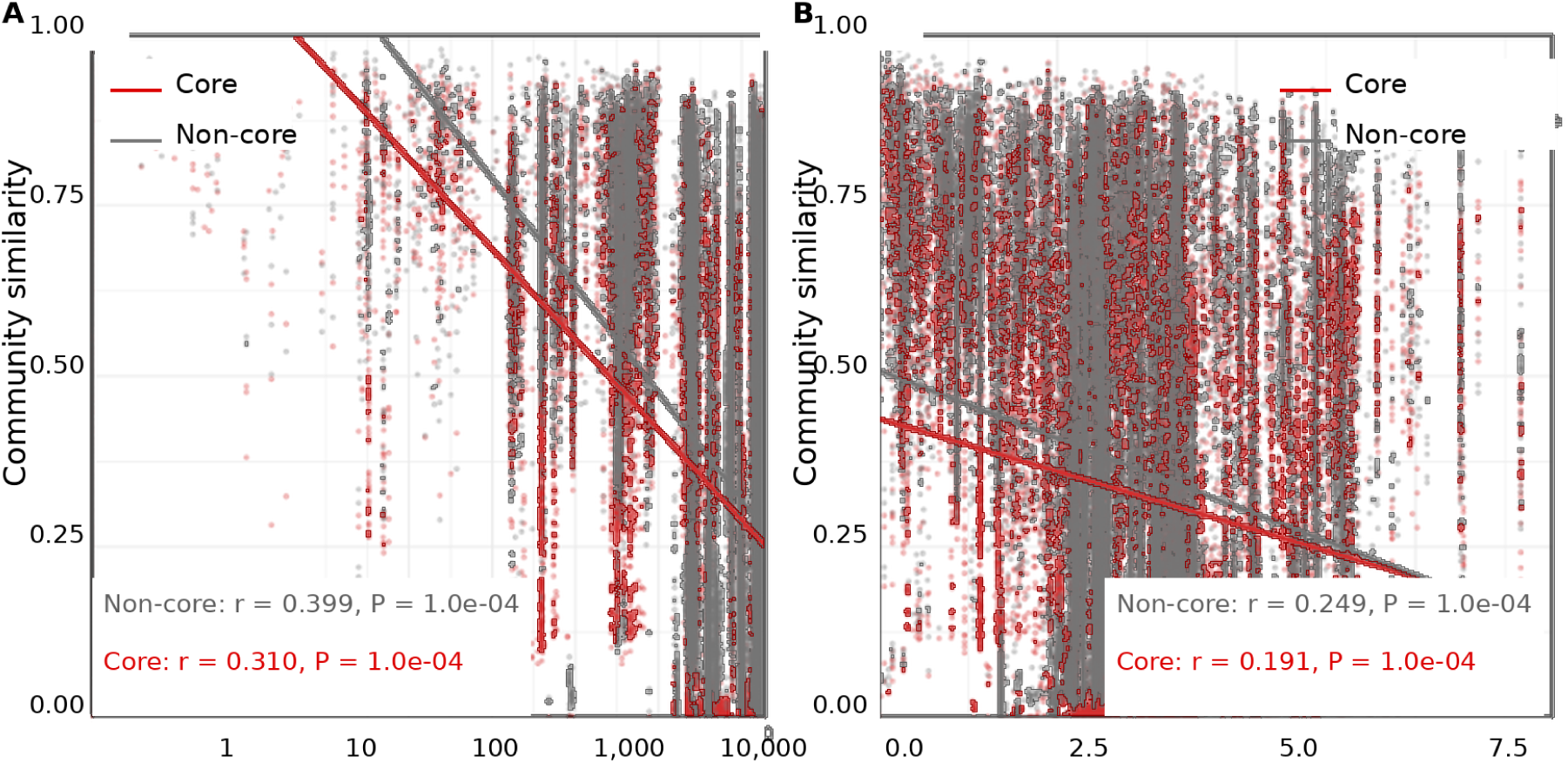
Spatial and environmental structuring of core and non-core microbial communities. (**A**) Distance-decay relationships showing pairwise community dissimilarity (Bray–Curtis) as a function of geographic distance (log₁₀ km) for core (red) and non-core (gray) fractions. Solid lines indicate linear regressions for each group. Both fractions showed significant distance-decay patterns (core: r = 0.310, P = 1.0e-4; non-core: r = 0,399, P = 1.0e-4), with the non-core community exhibiting a steeper decline in similarity with distance. (**B**) Environmental decay relationships showing community dissimilarity as a function of environmental distance. Linear fits are shown for core and non-core fractions. Both fractions exhibited significant correlations with environmental gradients (core: r = 0,191, P = 1.0e-04; non-core: r = 0.249, P = 1.0e-04), indicating strong environmental filtering across sampling sites.

Environmental drivers further accentuated this contrast. Variation in the non-core fraction was strongly associated with climatic and edaphic gradients, including precipitation regimes, temperature variability, and sediment properties. These taxa responded sensitively to local environmental conditions, leading to pronounced regional differentiation. By comparison, the core fraction exhibited limited responses to these gradients, consistent with broad environmental tolerance and stabilizing selection.

The disproportionate contribution of non-core taxa to biogeographic structure was evident across analytical approaches. Ordination analyses based on non-core taxa recapitulated the major east-west and regional patterns observed in the full dataset, whereas ordinations restricted to core taxa showed weak clustering by region or environment. Together, these results indicate that most spatial and environmental differentiation in mangrove-sediment microbiomes resides in the peripheral microbial community, consistent with core-satellite structures observed across microbial ecosystems^17,18,23^.

These contrasting patterns support a dual architecture in which a conserved microbial core provides functional continuity across mangrove ecosystems, while a large and dynamic periphery enables local adaptation and regional diversification. This organization reconciles the coexistence of global similarity in dominant microbial functions with strong biogeographic differentiation in community composition.

### Core taxa form the structural backbone of microbial networks

Co-occurrence network analyses revealed strong differences in the structural roles of core and non-core taxa within mangrove-sediment microbiomes. Networks constructed from the full dataset displayed highly connected architectures characterized by a small number of centrally positioned nodes. These nodes were disproportionately composed of core taxa, which consistently exhibited high degree, betweenness, and closeness centrality across regions (Fig. 6A), consistent with the central roles of core or hub taxa reported in microbial network analyses ^29,30^.

**Figure 6.**
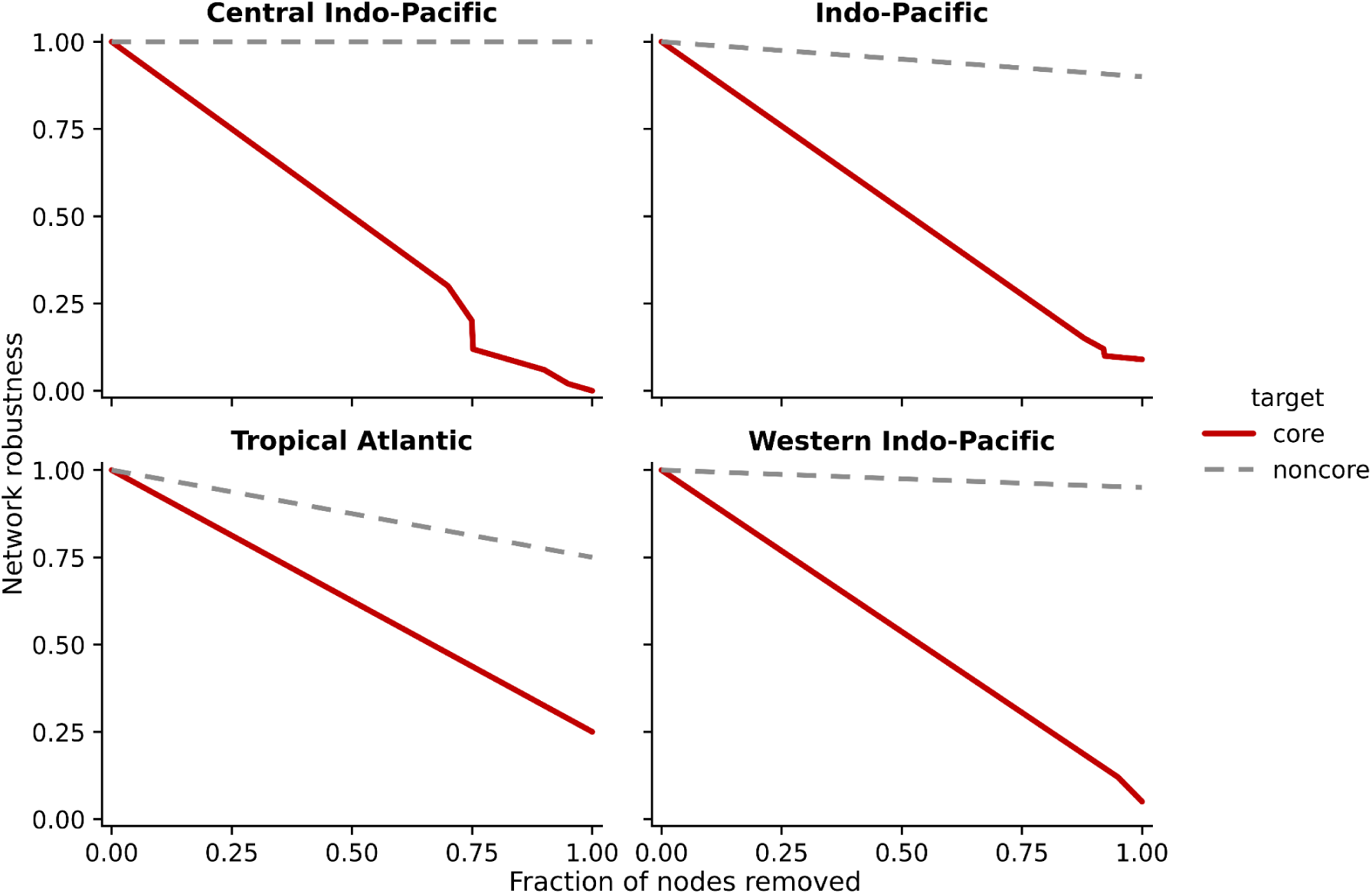
Network robustness under targeted removal of core vs non-core taxa. Attacks on high-degree core taxa (red solid lines) caused a rapid loss of connectivity and network collapse after removal of 50–65% of targeted core nodes, whereas removal of non-core taxa (grey dashed lines) left the main connected subnetwork largely intact even when most non-core nodes were removed. These patterns indicate that core taxa disproportionately occupy high-degree positions and form the structural backbone that stabilizes community associations across realms.

To assess the structural importance of core taxa, we performed targeted node-removal analyses. Removing core taxa resulted in rapid declines in network connectivity and fragmentation into smaller, poorly connected components. In contrast, random removal of non-core taxa produced comparatively minor changes in network structure. These patterns indicate that core taxa occupy positions critical for maintaining the overall topology of microbial co-occurrence networks ^29,30^

Importantly, the centrality of core taxa was consistent across marine realms and environmental contexts, despite pronounced regional differences in peripheral community composition. This consistency suggests that the structural role of core taxa reflects conserved ecological interactions associated with life in reduced mangrove sediments, rather than region-specific associations.

Together, these results show that mangrove-sediment microbial networks are organized around a conserved set of highly connected taxa that form a structural backbone, while a diverse and environmentally responsive periphery contributes most of the system’s compositional variability. This network organization provides an additional layer of evidence supporting the dual architecture of mangrove sediment microbiomes.

## CONCLUSION

We show that mangrove sediment microbiomes are organized around a dual architecture composed of a small, globally conserved core and a large, environmentally responsive periphery. The core persists across continents and climatic regimes, dominates community abundance and network connectivity, and is shaped primarily by deterministic selection. In contrast, non-core taxa respond strongly to climatic, edaphic, and macroecological gradients and account for most biogeographic turnover.

This organization explains how mangrove sediments maintain functional stability despite strong regional differentiation. By distinguishing a stable microbial backbone from a flexible peripheral community, our results provide a framework for predicting microbiome responses to environmental change and for incorporating microbial processes into mangrove monitoring, conservation, and restoration strategies.

## METHODS

### Metagenomic data acquisition, preprocessing, and taxonomic profiling

A total of 390 metagenomes from mangrove sediments were compiled from 43 sites across 12 countries within tropical and subtropical regions, generated between 2011 and 2021. Data were retrieved from the public repositories NCBI Sequence Read Archive (SRA) and European Nucleotide Archive (ENA). Files were preferentially downloaded in FASTQ format using prefetch and fasterq-dump from sra-tools v3.1.1; in cases where only SRA files were available, native conversion was performed. Geographical coordinates, sampling year, and associated metadata were recovered from each project’s supplementary information. Inclusion criteria were: environmental samples classified as “mangrove sediment”, shotgun metagenomes (non-amplicon). In addition, metadata were manually checked to ensure the provenance of mangrove sediment environments.

Quality control was performed with FastQC v0.11.9, and read trimming with fastp v0.23.4 under standard thresholds: automatic adapter trimming, Q ≥ 30, minimum length 50 bp, and removal of reads containing > 5% undefined bases. Taxonomic classification was conducted with Kraken2 v2.1.2 using the PlusPF database (complete RefSeq, January 2023 update). For downstream community analyses, OTU tables were derived and stored together with matching sample metadata.

### Environmental, anthropogenic, and biodiversity covariates

Climatic variables were extracted from open global datasets: WorldClim v2.1 (temperature and precipitation bioclimatic variables, 1 km² resolution), and the Human Footprint Index 2020 (anthropogenic pressure). Biological diversity variables were derived from GBIF (Global Biodiversity Information Facility) as estimates of species richness and Shannon diversity within 1 km buffers around sampling sites, corrected for sampling effort via spatial rarefaction.

Soil properties were obtained from SoilGrids v2.0, including organic carbon content (%C), total nitrogen (%N), pH in water, cation exchange capacity (CEC), bulk density (BD), soil texture (clay, silt, and sand fractions), and electrical conductivity (EC), standardized at 0-30 cm depth. All layers were reprojected to a common geographic system (WGS84), homogenized to 1 km² spatial resolution, and extracted per sampling coordinate using bilinear interpolation. Continuous variables were z-standardized before multivariate analyses.

### Alpha diversity and spatial modelling

#### Alpha diversity metrics

For alpha-diversity analyses, the OTU-by-sample count matrix was rarefied without replacement to the minimum sequencing depth across retained samples, to homogenize sequencing effort. From the rarefied matrix, we calculated the Hill numbers q = 0, 1, and 2. The primary metric was Hill q = 1, defined as the exponential of Shannon diversity. These values were integrated with the sample metadata and summarized by marine biogeographic realm (realm) using violin/box plots of Hill q = 1 and realm-level summary statistics (Supplementary Fig. S1; Supplementary Table S1).

### Environmental and spatial drivers of alpha diversity

To model the variation in microbial alpha diversity, we used generalized additive models (GAMs) fitted to Hill q = 1. Shannon diversity was computed from the OTU relative-abundance table (natural logarithm) and converted to Hill q = 1. For modelling, we used a log-transformed version of Hill q = 1, obtained by adding a constant of 1 to the diversity values and then taking the natural logarithm. This transformation stabilises variance and prevents problems with zero values.

As candidate predictors, we considered a set of non-collinear environmental and geographic covariates: climatic variables (BIO1-BIO19), human footprint, local GBIF-based richness and Shannon diversity, soil properties (including sand_mean, clay_mean, bdod_mean, soc_mean, phh2o_mean, silt_mean), and spatial coordinates (latitude, longitude). To reduce multicollinearity, we computed the Pearson correlation matrix among candidate predictors (pairwise complete observations) and removed variables with r≥ 0.90, retaining a non-collinear subset for modelling.

GAMs were fitted with the mgcv package using bam with Gaussian error and identity link. For each continuous predictor, we included a thin-plate regression spline s(x, bs = "tp") with basis dimension k chosen adaptively between 3 and 6 according to the number of unique observations. We included site as a random effect via s(Locality, bs = "re") to account for shared variation among samples from the same site. Models were estimated by fast restricted maximum likelihood (method = "fREML") with a modest penalty inflation (gamma = 1.4) to limit overfitting.

We first fitted a “full” model containing smooth terms for all predictors retained after the collinearity screen, plus the random-effect term. We then fitted a parsimonious “RF-based” model including only the subset of environmental variables selected in a preceding random forest analysis and present in the metadata (clay_mean, bdod_mean, BIO13, BIO3, BIO15, BIO2, gbif_shannon, and the locality random effect). Models were compared using Akaike’s information criterion (AIC), deviance explained, and adjusted R². Residual diagnostics (gam.check, residual-vs-fitted plots, Q-Q plots, and histograms) were inspected to assess distributional assumptions and leverage (Supplementary Table S2). For graphical presentation of smooth effects, fitted values and standard errors were first obtained on the log-transformed diversity scale and then converted back to the original Hill q = 1 scale. The resulting values were plotted as predicted means with 95% confidence bands.

Spatial structure in alpha diversity was further captured by including a two-dimensional smooth over latitude and longitude. Specifically, we used a tensor-product smooth te(lat, lon) to generate predicted maps of Hill q = 1 while holding non-spatial covariates at their median values. Predicted values over a latitude-longitude grid were visualized as spatial “hotspots” of diversity, overlaid with observed Hill q = 1 values and with country-level sampling intensity (Fig. 1A).

### Beta-diversity metrics and distance-decay relationships

We quantified distance-decay relationships for taxonomic community composition in mangrove-sediment samples using geographic distance and two dissimilarity indices: Bray-Curtis (relative abundance) and Sørensen (presence-absence). Community tables were filtered to remove taxa with zero total counts and samples lacking geographic coordinates. Bray-Curtis dissimilarities were computed on row-wise normalized abundance matrices (relative abundances), whereas Sørensen dissimilarities were obtained as binary Bray-Curtis distances computed on presence-absence data (vegdist, method = "bray", binary = TRUE, R package vegan).

Geographic distances between sampling sites were calculated from latitude-longitude coordinates using the function distm in the R package geosphere (v1.5-18) with the distHaversine option, which estimates great-circle distances on the WGS84 ellipsoid, and expressed in kilometres. To test whether community dissimilarity increased with geographic distance, we performed Mantel tests (Pearson correlation, 9,999 permutations) between each community dissimilarity matrix (Bray-Curtis or Sørensen) and the corresponding geographic distance matrix.

For visualization (Fig. 2A), dissimilarities were converted to similarities (1 − dissimilarity) and plotted against log₁₀-transformed geographic distances with linear fits. To compare the strength of distance-decay relationships between abundance-based and presence-absence metrics, we fitted linear models of community similarity as a function of log₁₀ distance and dissimilarity type (Bray-Curtis versus Sørensen), including their interaction: similarity ∼ log₁₀(distance) × metric. Differences in regression slopes were evaluated using a permutational analysis of covariance (ANCOVA), in which the F statistic for the interaction term was compared to a null distribution generated by randomly permuting the dissimilarity-type labels (999 permutations).

### Ordination and regional structure

Non-metric multidimensional scaling (NMDS) was performed on taxonomic community tables filtered to mangrove-sediment samples. For the Bray-Curtis analysis, OTU counts were first converted to relative abundances at the sample level, and Bray-Curtis dissimilarities were computed from this table. In parallel, compositional differences were quantified using Aitchison distances, obtained from centred log-ratio (CLR) transformed compositions after adding a small pseudocount and renormalizing sample totals; Euclidean distances in CLR space were used as Aitchison distances.

NMDS ordinations (k = 2) were obtained with metaMDS (vegan) from the Bray-Curtis and Aitchison distance matrices, respectively. Site scores from the Aitchison-based NMDS were subsequently used to fit smooth environmental surfaces for geographic longitude using generalized additive models (GAMs; mgcv, model z ∼ s(NMDS1, NMDS2), REML). Predicted values over a regular grid in NMDS space were visualized as isolines overlaid on sample scores coloured by realm (Fig. 2B).

Biogeographic factors (ecoregion, province, realm, and country) were represented by different numbers of sites. To minimize biases caused by unbalanced sampling when testing for compositional differences among regions, we complemented PERMANOVA analyses with a balanced subsampling approach based on the same relative-abundance table used for the Bray-Curtis NMDS. For each factor, we first excluded levels represented by fewer than n sites (here n = 5). We then identified the minimum sample size across the remaining levels (n_min) and repeatedly drew balanced subsets by randomly selecting n_min sites per level without replacement. For each balanced subset, we recomputed Bray-Curtis dissimilarities and performed PERMANOVA (adonis2, vegan) with 999 permutations. This procedure was iterated 199 times per factor, yielding a distribution of F statistics, R², and permutation-based P values. The robustness of regional effects to uneven sampling effort was assessed by inspecting the distribution of F, R², and P across iterations; results are summarized in Supplementary Table S3 and Supplementary Fig. S2.

### Variance partitioning of beta diversity

We quantified how environmental and spatial factors structure mangrove-sediment community composition using distance-based redundancy analysis (db-RDA) on Bray-Curtis dissimilarities computed from the OTU table. Predictors were organized into four sets: (i) a large-scale geographic gradient (linear latitude and longitude), (ii) space among realms (categorical realm), (iii) space within realms represented by Moran’s eigenvector maps (MEM/dbMEM), and (iv) environment, the same non-collinear set of predictors used in the full GAM (climate, human footprint, GBIF metrics, and soil properties). MEMs are orthogonal spatial eigenfunctions derived from geographic distances that capture spatial autocorrelation at multiple spatial scales. Here, MEMs were computed separately within each realm from Euclidean distances on coordinates and retained if non-constant. Including MEMs allows the model to absorb fine-to meso-scale spatial structure that often co-varies with the environment, preventing inflation of “pure environment” effects.

We fit two complementary specifications. The BASE model included all predictor sets (ENV + lat/lon + realm + MEM) and thus partitions variance into (i) pure effects of each set and (ii) a shared environment-space component reflecting their co-distributed structure. To assess how much apparent environmental signal is attributable to unmodelled spatial autocorrelation, we also fit a NO-MEM model (ENV + lat/lon + realm) that omits within-realm MEMs; this provides an upper bound for the “pure environment” fraction when fine-scale space is not explicitly modelled. For each specification, we report the adjusted R² of the global db-RDA and obtain pure fractions via partial db-RDAs that condition on the remaining sets. The shared environment-space fraction is computed as R² global minus the sum of pure fractions; residual variance equals 1 - R² global. Significance of the global model and of each pure fraction was assessed with 999 permutations. A db-RDA biplot from the environment-only constrained ordination (sites coloured by realm with the most significant envfit vectors) is provided to aid interpretation of dominant gradients

### Community assembly processes

Community assembly processes were quantified using iCAMP, which partitions taxonomic turnover into discrete ecological processes based on phylogenetic binning and null-model comparisons. For this analysis, we used a phyloseq object in which metagenomic taxa had been aggregated at the genus level and linked to a genus-level phylogenetic tree. Only samples annotated as “mangrove sediment” in the EcologicalClass field were retained, matching the subset used in the diversity and beta-diversity analyses.

The resulting genus-by-sample matrix (390 samples × 500 genera) and the corresponding phylogenetic tree were passed to the icamp.cm function (R package iCAMP) using Bray-Curtis dissimilarity and the β mean nearest taxon distance (bMNTD) as the phylogenetic metric. Marine realm (realm) was used as the meta.group factor so that null models were stratified by realm. To reduce computational burden while retaining resolution at the genus level, the analysis was restricted to the 500 most abundant genera across samples (N_TOP_GENUS = 500).

For each pair of communities, iCAMP decomposes compositional turnover into the relative contributions of five processes: heterogeneous selection (HeS), homogeneous selection (HoS), dispersal limitation (DL), homogenizing dispersal (HD), and drift/weak selection (DR), following the framework of Ning et al. Null expectations were generated using the Raup-Crick (RC) index with bMNTD and 999 randomizations and corrected for special cases as implemented in icamp.cm. Bins were defined using a phylogenetic-bin size limit of 24 taxa (bin.size.limit = 24).

The icamp.cm output was summarized at the community-pair level into a table with the relative importance (proportions summing to 1) of HeS, HoS, DL, HD, and DR for each of the 75,855 pairwise comparisons among the 390 mangrove-sediment communities. We then (i) averaged these proportions across all pairwise comparisons to obtain the mean contribution of each process and (ii) assigned each community pair to a “dominant” process defined as the one with the highest relative importance. These summaries are reported in Supplementary Fig. S3 and Supplementary Table S4.

### Core microbiome components

Technical and biological replicates from the same site were aggregated by summing OTU counts per Locality, yielding an OTU-by-locality matrix. To minimise the influence of sequencing artefacts and extremely rare variants, we first applied a global prevalence filter, retaining only OTUs with ≥ 50 total reads across all mangrove localities and detected in at least 5 localities.

For each retained OTU, we calculated (i) its prevalence and mean relative abundance within each biogeographic realm (realm), based on the locality-level matrix, and (ii) its distribution across localities. An OTU was classified as core if, in at least one realm, it occurred in ≥ 40% of localities (prevalence ≥ 0.4) and its mean relative abundance in that realm was ≥ 0.01% (1×10⁻⁴). OTUs detected in exactly one realm were defined as specific. To quantify spatial concentration, we computed locality dominance for each OTU, defined as the fraction of its total relative abundance across all localities that occurred in the locality with the highest relative abundance; OTUs with locality dominance ≥ 0.8 were labelled indicators. OTUs present in a single locality only were referred to as unique.

These binary properties were combined into mutually exclusive components defined as follows: (i) core and indicator, OTUs classified as both core and indicator; (ii) core, OTUs classified as core but not indicator; (iii) specific and indicator, OTUs specific to a single realm and also indicator, but not core; (iv) indicator, OTUs classified as indicator but neither core nor specific; (v) specific and unique, OTUs specific to a single realm and occurring in a single locality only; and (vi) specific, OTUs specific to a single realm but neither indicator, unique nor core (Supplementary Fig. S4). OTUs occurring in more than one realm that did not meet the criteria for core, specific, or indicator were classified as multi-realm generalists, and any remaining OTUs were assigned to other. Using these thresholds on the filtered dataset, 21,443 OTUs were classified into: core (819, 3.8%), core and indicator (50, 0.2%), indicator (447, 2.1%), specific and indicator (1, < 0.01%), specific and unique (0), specific (1, < 0.01%) and multi-realm generalists (20,125, 93.9%); no OTUs were assigned to the other category. These components are summarised in Fig. 4A and listed in full, together with taxonomic annotations, in Supplementary Data S1.

To characterise rare realm-restricted taxa that are removed by this global prevalence filter, we additionally quantified, for each realm, the number of OTUs that were detected exclusively in that realm and the subset of those that appeared in a single locality only. This realm-specific analysis was performed on the unfiltered OTU table (i.e., including all OTUs with at least one read) and is shown in Fig. 4B.

### Distance-decay and environment-decay patterns: Core vs. non-core microbiome

We quantified distance-decay and environment-decay relationships separately for core and non-core components of the mangrove-sediment microbiome. Microbial community analyses were conducted in R (v4.4.0) using the phyloseq package for microbiome data integration (phyloseq v1.50.0) and vegan for ecological distance calculations (vegan v2.7-1). Core and non-core OTUs were classified based on previously defined component categories (“core”, “core and indicator”, “indicator”, “multi-realm generalist”, “specific”, and “specific and indicator”). Community dissimilarities were quantified using Bray–Curtis distances computed on OTU tables separated into core and non-core subsets. Phyloseq objects were filtered to retain only mangrove sediment samples (EcologicalClass = "Mangrove sediment") and further subset to create separate community matrices for core-only and non-core-only OTUs. Samples with zero total abundance after subsetting were removed. To ensure consistent pairwise comparisons, only samples present in both core and non-core datasets and possessing complete geographic coordinates and environmental data were retained for downstream analyses (n = 329 samples).

Bray-Curtis dissimilarities were computed on row-wise normalized abundance matrices (relative abundances) for both core and non-core communities using the vegdist function (method = "bray") in the R package vegan (v2.7-1). Geographic distances between sampling sites were calculated from latitude-longitude coordinates using the distm function in the R package geosphere (v1.5-18) with the distHaversine option, which estimates great-circle distances on the WGS84 ellipsoid, and expressed in kilometres. Environmental distances were calculated as Euclidean distances among scaled environmental variables (bio1, bio3, bio13, bio15, gbif_shannon_H_2km, gbif_richness_species_2km, sand_mean) using the dist function.

To test whether community dissimilarity increased with geographic and environmental distance, we performed Mantel tests (Pearson correlation, 9,999 permutations) between each community dissimilarity matrix (core or non-core) and the corresponding distance matrix (geographic or environmental) using the mantel function in vegan. To quantify and compare the rates of compositional turnover, we fitted linear regression models relating community similarity (1 - Bray-Curtis dissimilarity) to log₁₀-transformed geographic distance and to environmental distance separately for core and non-core communities. We then used analysis of covariance (ANCOVA) to test whether the slopes of these relationships differed significantly between core and non-core fractions, with community type (core vs. non-core) as a categorical predictor and its interaction with distance as the focal term. All statistical analyses were conducted in R (v4.4.0).

### Environmental drivers of core richness and dominance

To test whether the climatic thresholds detected for overall diversity also structure the mangrove core microbiome, we extended the generalized additive model (GAM) framework used for alpha diversity to three related responses: (i) total OTU richness, (ii) richness of core OTUs and (iii) the fraction of sequencing reads assigned to core OTUs per sample (“core dominance”). We computed for each sample: total richness (number of OTUs with > 0 reads), core richness (number of OTUs assigned to the core or core-and-indicator components), non-core richness (all remaining OTUs), and the fraction of reads belonging to core OTUs (frac_reads_core). Richness variables were transformed as log(1 + richness); core dominance was transformed with a logit after clipping fractions to [10⁻⁴, 1 - 10⁻⁴] to avoid extreme values. For comparability with the alpha-diversity GAMs, all models used the same reduced set of key environmental predictors. Locality was included as a random effect via s(Locality, bs = "re") to account for shared variation among samples from the same site.

Each response was modelled with mgcv::bam assuming Gaussian error and identity link, using thin-plate regression splines s(x, bs = "tp") with basis dimension k between 3 and 6, chosen according to the number of unique observations. Models were estimated by fast restricted maximum likelihood (method = "fREML", gamma = 1.4), as in the alpha-diversity GAMs. Fitted values and standard errors were obtained on the transformed scale (excluding the random term) and then back-transformed for plotting. Partial effects were visualised as smooth curves with 95% confidence bands for each predictor (Supplementary Fig. S5; Supplementary Table S5).

### Realm-specific backbone networks and robustness to node removal

To evaluate the structural role of the core taxonomic fraction, we built separate co-occurrence networks for each marine realm using the same OTU-by-locality matrix and core-component labels defined in the core-microbiome analysis. Within each realm, technical replicates were aggregated by summing OTU counts per locality. We then applied prevalence and abundance filters (minimum number of localities and total reads per OTU) to restrict the network to robustly detected taxa and reduce the influence of extremely rare OTUs on association estimates. Abundance tables were converted to relative abundances, and pairwise Spearman correlations were computed among OTUs. For each realm, we constructed an undirected co-occurrence network retaining only OTU pairs with ρ ≥ 0.6 and false-discovery-rate-corrected significance; self-loops were removed.

For downstream analyses, we focused on the main connected subnetwork, defined as the largest set of nodes in which every node can be reached from any other through at least one path. For this main subnetwork, we recorded the number of nodes and edges, mean degree (2E/N), and the fraction of nodes annotated as core (i.e., OTUs belonging to the core or core-and-indicator components in the global classification; all remaining OTUs were labelled non-core).

To quantify how core and non-core taxa contribute to network robustness, we simulated targeted attacks on each realm-specific network. Nodes were ordered by degree (from highest to lowest), and two separate target sets were defined: (i) core nodes only and (ii) non-core nodes only. For each target set, nodes were removed sequentially following this degree order, while nodes in the other set were kept intact. After each removal step, we recomputed the size of the main connected subnetwork and expressed it as a fraction of its initial size (relative main-subnetwork size). The progress of the attack was expressed as the fraction of targeted nodes removed.

For each realm and target set, the resulting curves (relative main-subnetwork size vs. fraction of targeted nodes removed) were summarised by the area under the curve (AUC) estimated by trapezoidal integration on [0, 1]. Low AUC values indicate that the main subnetwork collapses quickly as nodes are removed (low robustness), whereas high AUC values indicate that connectivity is largely preserved throughout the attack. realm-specific network properties and robustness metrics are reported in Supplementary Table S6.

## FUNDING

Financial support for this study was provided by the Secretaría de Ciencia, Humanidades, Tecnología e Innovación (SECIHTI) through the Ciencia de Frontera program (Project CBF2023-2024-1251), and by the Universidad Nacional Autónoma de México (UNAM) through the Programa de Apoyo a Proyectos de Investigación e Innovación Tecnológica (PAPIIT) (Project IA201824). F.J.B.O. was supported by a SECIHTI postdoctoral fellowship (474433).

## Notes

### Competing Interest Statement

The authors have declared no competing interest.

