## Supplementary material for "Global biogeography of mangrove sediment microbiomes is structured by a conserved core and environmental selection"

**Supplementary materials**

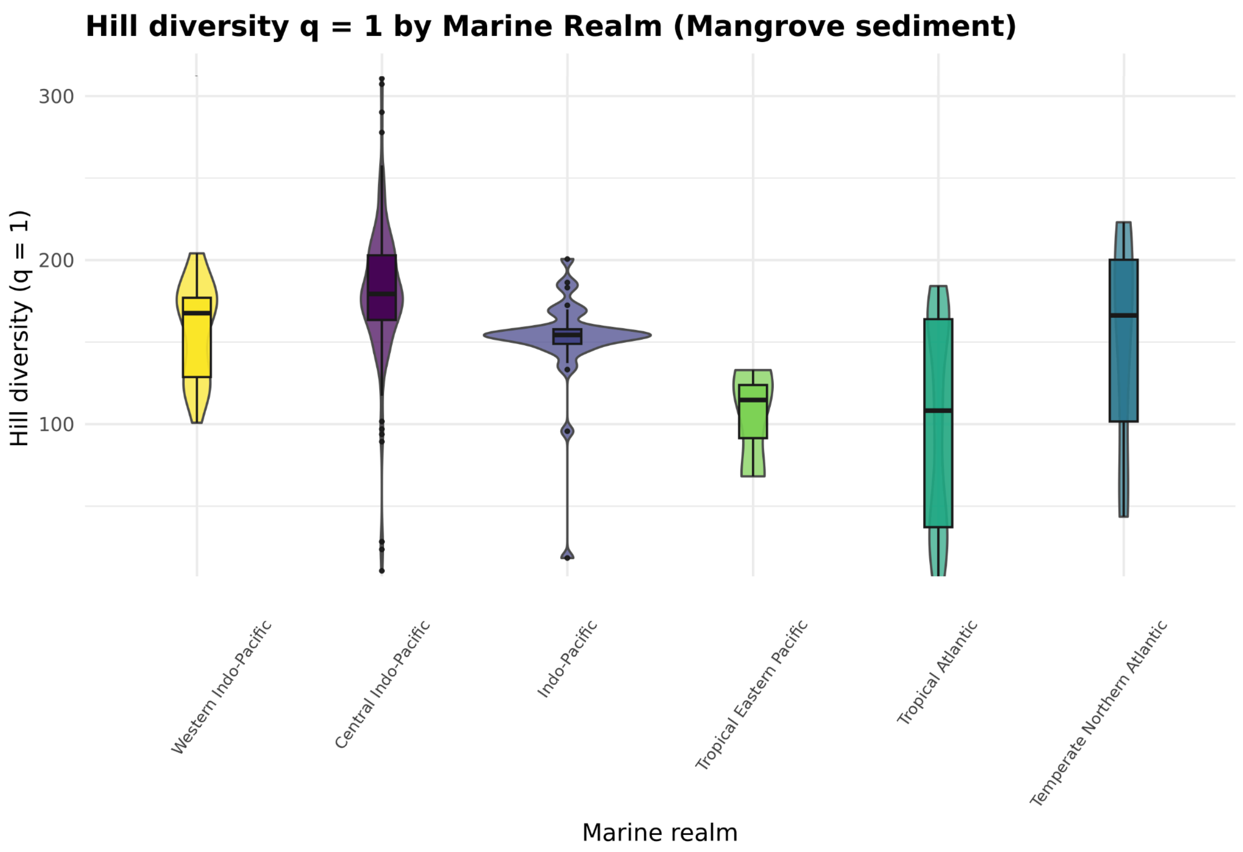


**Figure S1. Hill diversity (q = 1) across marine realms in mangrove sediment.** Distribution of Hill diversity of order 1 (q = 1; equivalent to the exponential of the Shannon index) for samples grouped by marine real. Violin plots depict the data density; boxplots show the median and interquartile range (IQR), with whiskers extending to 1.5×IQR; points indicate individual observations and outliers.

**Table S1.** Realm-level summary statistics of rarefied alpha diversity (Hill numbers q = 0, 1, and 2) in mangrove sediment samples.

| **Realm** | **n_samples** | **Hill q=1 mean ± SD** | **Hill q=1 median [IQR]** | **Hill q=1 range** | **Hill q=0 mean ± SD** | **Hill q=2 mean ± SD** |
|---|---|---|---|---|---|---|
| **Central Indo-Pacific** | 171 | 180.27 ± 42.06 | 179.40 [163.52–202.95] | 10.53–310.67 | 8959 ± 1434 | 25.98 ± 4.45 |
| **Indo-Pacific** | 33 | 151.38 ± 29.43 | 154.35 [148.93–157.79] | 18.35–200.64 | 8139 ± 1075 | 24.75 ± 4.66 |
| **Temperate Northern Atlantic** | 12 | 149.53 ± 65.32 | 166.30 [101.58–200.21] | 43.54–223.08 | 7932 ± 2417 | 25.29 ± 4.78 |

| | | | | | | |
|---|---|---|---|---|---|---|
| **Tropical Atlantic** | 139 | 100.12 ± 63.52 | 108.25 [37.15–163.99] | 5.92–184.19 | 4863 ± 3799 | 18.37 ± 7.78 |
| **Tropical Eastern Pacific** | 3 | 105.27 ± 33.42 | 114.73 [91.44–123.83] | 68.15–132.94 | 7529 ± 868 | 18.26 ± 3.86 |
| **Western Indo-Pacific** | 32 | 157.35 ± 29.03 | 167.63 [128.67–177.03] | 100.83–204.12 | 8729 ± 756 | 24.15 ± 3.06 |

**Table S2.** Smooth-term significance tests from the parsimonious GAM of log-transformed Hill q = 1 alpha diversity in mangrove sediments.

| Predictor | edf | Ref.df | F | p-value |
|---|---|---|---|---|
| **s(Human footprint 2020)** | 0.000 | 5 | 0.000 | 0.937 |
| **s(Annual mean temperature-BIO1)** | 0.772 | 5 | 0.951 | 0.006 |
| **s(Mean diurnal range-BIO2)** | 0.000 | 5 | 0.000 | 0.297 |
| **s(Isothermality-BIO3)** | 0.951 | 5 | 5.458 | 0.000 |
| **s(Precipitation of the wettest month-BIO13)** | 2.867 | 5 | 21.277 | 0.000 |
| **s(Precipitation seasonality-BIO15)** | 2.598 | 5 | 7.401 | 0.000 |
| **s(Regional diversity (Shannon)-GBIF)** | 0.648 | 5 | 0.515 | 0.025 |
| **s(Regional richness-GBIF species richness)** | 2.296 | 5 | 6.431 | 0.000 |
| **s(Soil organic carbon-SOC mean)** | 0.000 | 5 | 0.000 | 0.501 |
| **s(Soil pH ($H_2O$)-mean)** | 0.000 | 5 | 0.000 | 0.605 |
| **s(Soil sand content-mean)** | 0.889 | 5 | 2.249 | 0.000 |
| **s(Soil silt content-mean)** | 0.000 | 5 | 0.000 | 0.704 |
| **s(Soil clay content-mean)** | 0.000 | 5 | 0.000 | 0.278 |
| **s(Soil bulk density (oven-dry)-BDOD mean)** | 0.000 | 5 | 0.000 | 0.923 |
| **s(Geographic latitude)** | 0.000 | 5 | 0.000 | 0.885 |
| **s(Geographic longitude)** | 1.124 | 5 | 1.458 | 0.000 |
| **s(Sampling site-Locality random effect)** | 0.000 | 32 | 0.000 | 0.937 |

**Table S3.** Balanced-subsampling PERMANOVA results for Bray–Curtis community dissimilarities across biogeographic factors.

| Factor | Levels retained (>=5 sites) | Balanced n per level (n_min) | Iterations | F median [IQR] | $R^2$ median [IQR] | P median [IQR] | $P < 0.05$ (proportion) |
|---|---|---|---|---|---|---|---|
| **Country** | 9 | 5 | 199 | 3.646 [3.322–3.997] | 0.448 [0.425–0.470] | 0.001 [0.001–0.001] | **1** |
| **Ecoregion** | 9 | 8 | 199 | 6.061 [5.446–6.861] | 0.435 [0.409–0.466] | 0.001 [0.001–0.001] | **1** |
| **Province** | 9 | 6 | 199 | 3.426 [3.057–3.786] | 0.378 [0.352–0.402] | 0.001 [0.001–0.001] | **1** |
| **Realm** | 5 | 12 | 199 | 4.769 [4.264–5.274] | 0.258 [0.237–0.277] | 0.001 [0.001–0.001] | **1** |

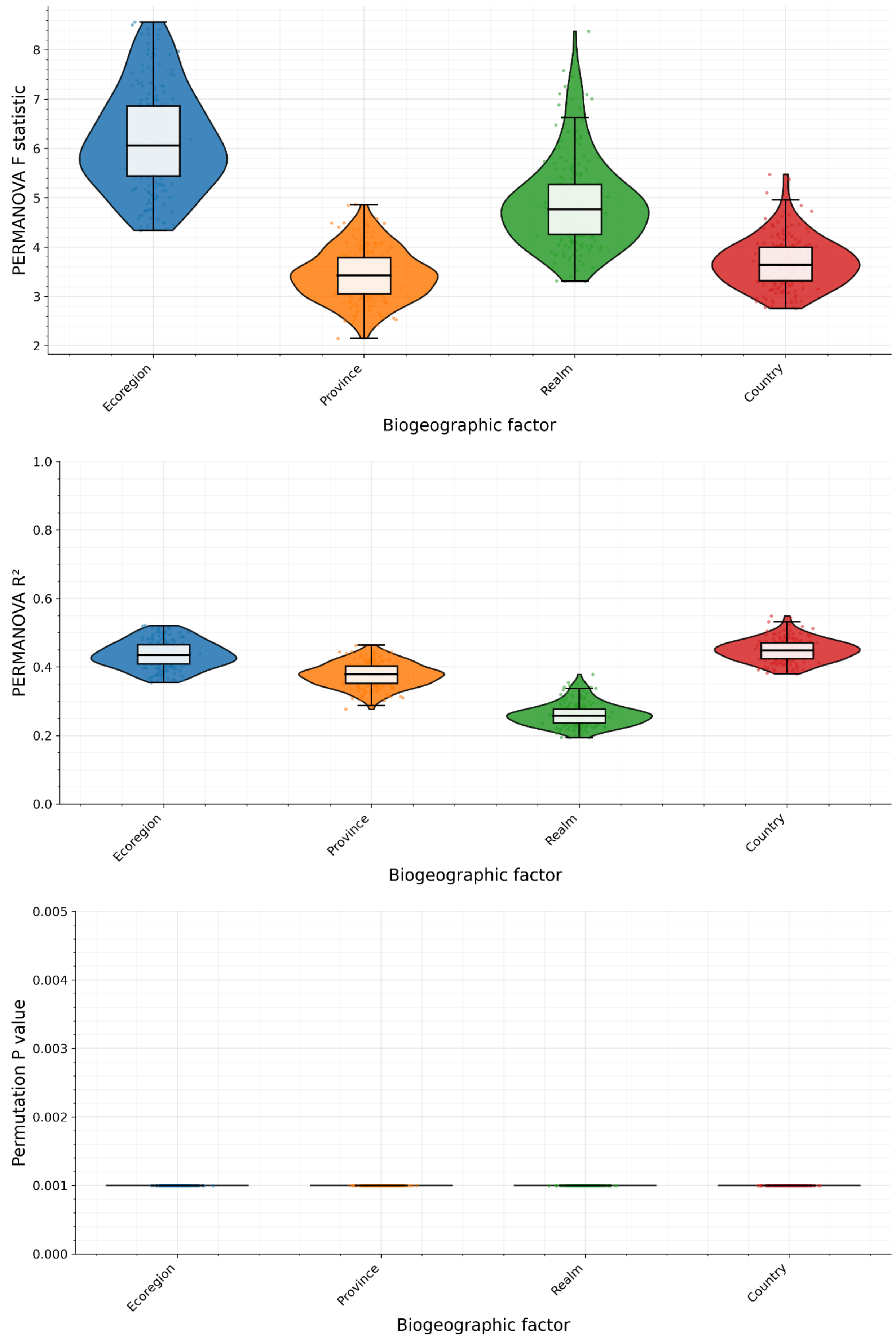
PERMANOVA F statistic
2
3
4
5
6
7
8
Ecoregion
Province
Realm
Country
Biogeographic factor
PERMANOVA R²
0.0
0.2
0.4
0.6
0.8
1.0
Ecoregion
Province
Realm
Country
Biogeographic factor
Permutation P value
0.000
0.001
0.002
0.003
0.004
0.005
Ecoregion
Province
Realm
Country
Biogeographic factor

**Figure S2. Robustness of regional structure under balanced subsampling PERMANOVA.** Distributions of PERMANOVA outputs across 199 balanced subsampling iterations for each biogeographic factor (Ecoregion, Province, Realm, and Country), after excluding levels represented by fewer than five sites and subsampling the remaining levels to the minimum sample size per level (n_min). For each iteration, Bray–Curtis dissimilarities were recomputed from sample-level relative abundances and tested with PERMANOVA (adonis2; 999 permutations). Boxplots summarize the iteration-wise F statistics (top), $R^2$ values (middle), and permutation-based P values (bottom). Across iterations, results were highly consistent for all factors P values remained uniformly low, and the distributions of F and $R^2$ were comparatively narrow, indicating that the detected regional structure in community composition is robust to uneven sampling effort and not driven by a small subset of overrepresented regions.

**Table S4.** iCAMP-based partitioning of community assembly processes from genus-level turnover among mangrove-sediment communities.

| **Process** | **Mean** | **CI_low** | **CI_high** | **Dominant_n** | **Dominant_%** |
|---|---|---|---|---|---|
| **Heterogeneous selection (HeS)** | 0.3 | 0.298 | 0.302 | 26545 | 34.994 |
| **Homogeneous selection (HoS)** | 0.272 | 0.27 | 0.274 | 24014 | 31.658 |
| **Dispersal limitation (DL)** | 0.295 | 0.294 | 0.297 | 19995 | 26.36 |
| **Homogenizing dispersal (HD)** | 0.113 | 0.112 | 0.114 | 5219 | 6.88 |
| **Drift / weak selection (DR)** | 0.02 | 0.019 | 0.02 | 82 | 0.108 |

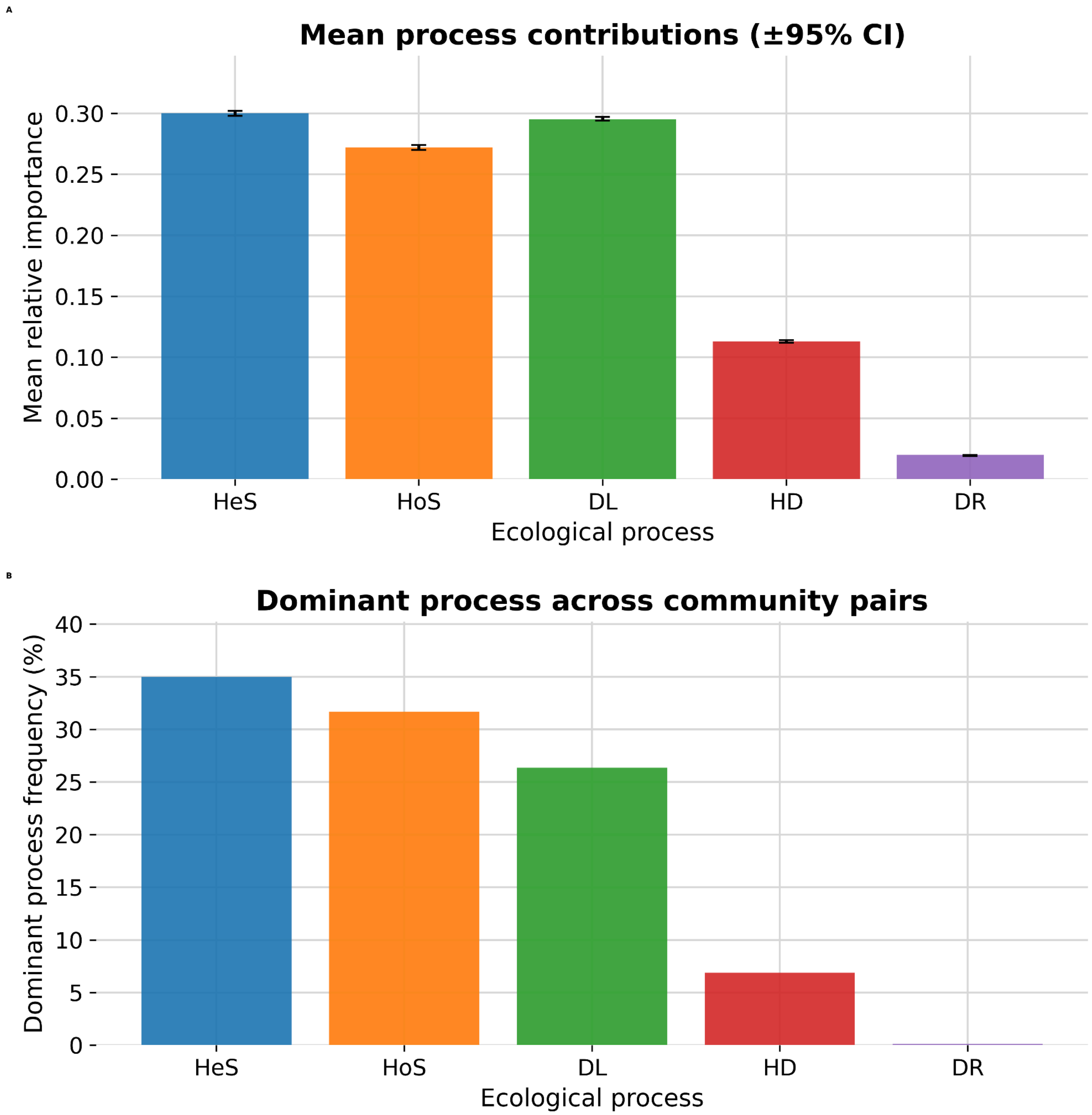


**Figure S3. Community assembly processes inferred by iCAMP across mangrove-sediment communities.** iCAMP was applied to genus-level community data (390 samples; 500 most abundant genera) using Bray–Curtis dissimilarities and bMNTD with 999 randomizations (bin.size.limit = 24), producing process contributions for each of 75,855 pairwise community comparisons. (**A**) Mean relative importance (proportion) of heterogeneous selection (HeS), homogeneous selection (HoS), dispersal limitation (DL), homogenizing dispersal (HD), and drift/weak selection (DR); error bars show 95% confidence intervals across all community pairs. (**B**) Frequency with which each process was the dominant contributor (largest proportion) across community pairs. Overall, turnover is primarily attributed to selection and dispersal-related processes (HeS, HoS, and DL), whereas homogenizing dispersal contributes less, and drift/weak selection is rare.

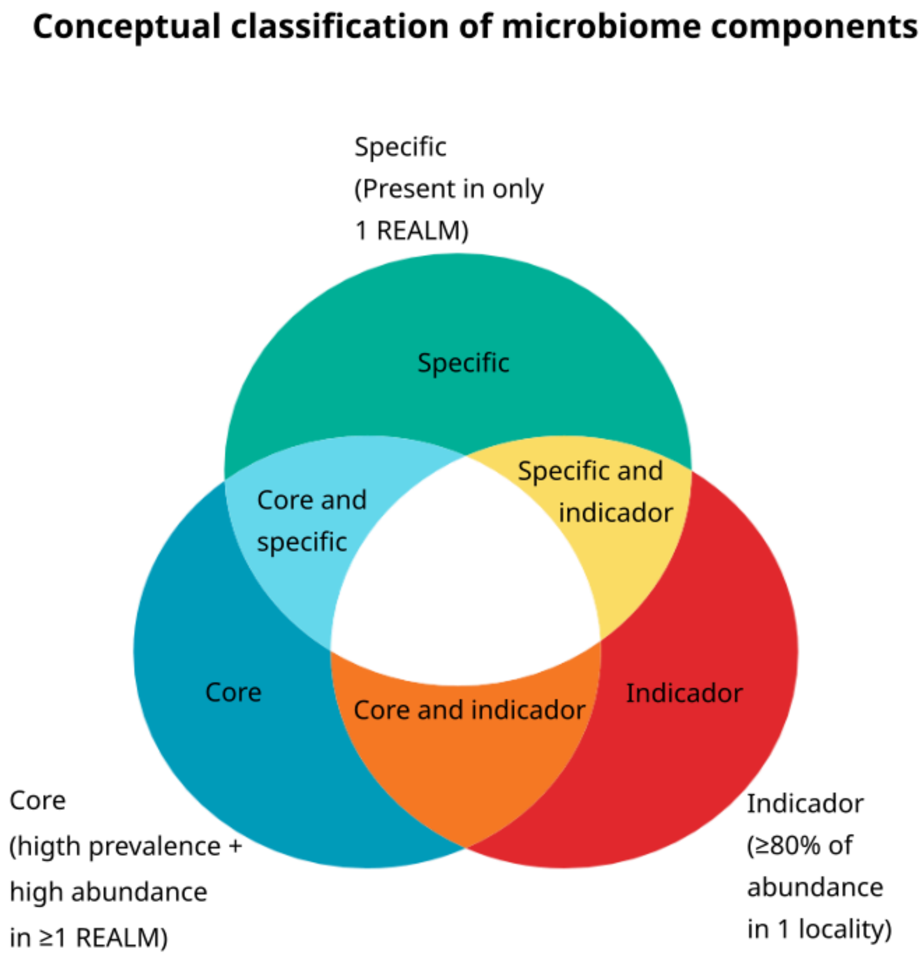


**Figure S4. Conceptual classification of microbiome components used to define core, specific, and indicator OTUs.** Schematic overview of the binary criteria applied to locality-aggregated OTU data to assign OTUs to microbiome components. OTUs were labelled core if, in at least one marine realm, they occurred in ≥40% of localities (prevalence ≥0.4) and had a mean relative abundance ≥0.01% ($1\times10^{-4}$) within that realm. OTUs detected in exactly one realm were labelled specific. OTUs were labelled indicators when strongly spatially concentrated, defined by locality dominance ≥0.8 (i.e., ≥80% of total relative abundance across all localities occurring in the single locality with maximum abundance). OTUs present in a single locality only were considered unique. Overlaps among these criteria yield mutually exclusive components (e.g., core and indicator; specific and indicator), which were used for downstream summaries.

**Table S5.** GAM smooth-term tests for environmental predictors of core dominance in mangrove-sediment communities.

| Predictor (smooth term) | edf | Ref.df | F | p_value |
|---|---|---|---|---|
| **s(Annual mean temperature-BIO1)** | 0.826 | 5 | 1.327 | 0.002 |
| **s(Isothermality-BIO3)** | 1.193 | 5 | 6.03 | 0 |
| **s(Precipitation of the wettest month-BIO13)** | 1.456 | 5 | 7.282 | 0 |

| | | | | |
|---|---|---|---|---|
| **s(Precipitation seasonality-BIO15)** | 2.361 | 5 | 7.375 | 0 |
| **s(Regional diversity (Shannon)-GBIF, 2 km)** | 0 | 5 | 0 | 0.484 |
| **s(Regional richness-GBIF species richness)** | 1.971 | 5 | 3.755 | 0 |
| **s(Soil sand content-mean)** | 0 | 5 | 0 | 0.364 |
| **s(Sampling site-Locality random effect)** | 0 | 32 | 0 | 0.757 |

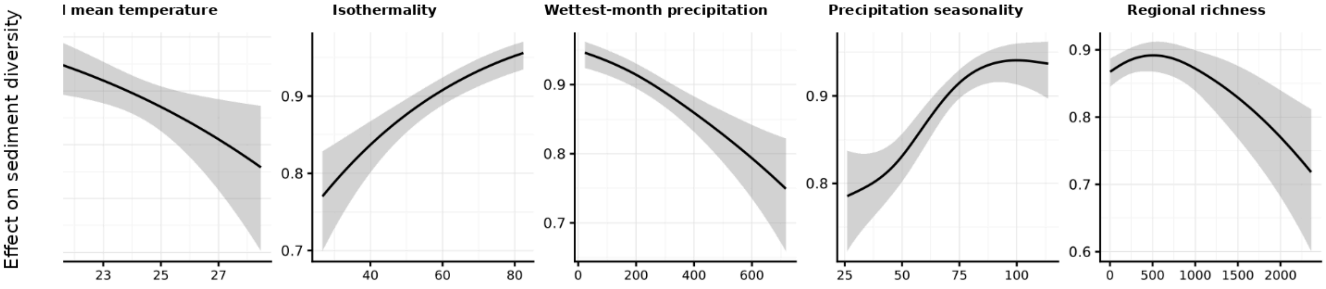


**Figure S5. Significant environmental drivers of core dominance in mangrove sediments.** Partial smooth effects from a generalized additive model (mgcv::bam; Gaussian error, identity link) fitted to logit-transformed core dominance (fraction of sequencing reads assigned to core OTUs per sample). Core OTUs were defined as those classified as Core or Core and indicator in the core-component framework. Smooth terms (thin-plate regression splines) were estimated with fREML and a penalty inflation (gamma = 1.4), including locality as a random effect (not shown). Only predictors with significant smooth terms ($p < 0.05$) are displayed. Black lines show fitted partial effects on the response scale, with grey ribbons indicating 95% confidence bands (±2 SE); rug marks along the x-axes indicate the distribution of observed predictor values.

**Table S6**. Realm-specific co-occurrence network properties and robustness to targeted removal of core versus non-core nodes.

| Marine realm | Localities (n) | Taxa in network (after filters) | Nodes (N) | Edges (E) | Core nodes fraction | AUC (targeted core removal) | AUC (targeted non-core removal) | Mean degree (2E/N) |
|---|---|---|---|---|---|---|---|---|
| **Central Indo-Pacific** | 171 | 587 | 587 | 31955 | 0.998 | 0.485 | 0.999 | 108.88 |
| **Indo-Pacific** | 33 | 562 | 562 | 50339 | 0.911 | 0.543 | 0.956 | 179.14 |
| **Tropical Atlantic** | 139 | 612 | 612 | 51520 | 0.765 | 0.617 | 0.882 | 168.37 |

| | | | | | | | | |
|---|---|---|---|---|---|---|---|---|
| **Western Indo-Pacific** | 32 | 551 | 551 | 33617 | 0.956 | 0.519 | 0.978 | 122.02 |
